# PTEN knockout using retrogradely transported AAVs restores locomotor abilities in both acute and chronic spinal cord injury

**DOI:** 10.1101/2023.04.17.537179

**Authors:** Andrew N. Stewart, Reena Kumari, William M. Bailey, Ethan P. Glaser, Gabrielle V. Hammers, Olivia H. Wireman, John C. Gensel

## Abstract

Restoring function in chronic stages of spinal cord injury (SCI) has often been met with failure or reduced efficacy when regenerative strategies are delayed past the acute or sub-acute stages of injury. Restoring function in the chronically injured spinal cord remains a critical challenge. We found that a single injection of retrogradely transported adeno-associated viruses (AAVrg) to knockout the phosphatase and tensin homolog protein (PTEN) in chronic SCI can effectively target both damaged and spared axons and restore locomotor functions in near-complete injury models. AAVrg’s were injected to deliver cre recombinase and/or a red fluorescent protein (RFP) under the human Synapsin 1 promoter (hSyn1) into the spinal cords of C57BL/6 PTEN^FloxΔ^/^Δ^ mice to knockout PTEN (PTEN-KO) in a severe thoracic SCI crush model at both acute and chronic time points. PTEN-KO improved locomotor abilities in both acute and chronic SCI conditions over a 9-week period. Regardless of whether treatment was initiated at the time of injury (acute), or three months after SCI (chronic), mice with limited hindlimb joint movement gained hindlimb weight support after treatment. Interestingly, functional improvements were not sustained beyond 9 weeks coincident with a loss of RFP reporter-gene expression and a near-complete loss of treatment-associated functional recovery by 6 months post-treatment. Treatment effects were also specific to severely injured mice; animals with weight support at the time of treatment lost function over a 6-month period. Retrograde tracing with Fluorogold revealed viable neurons throughout the motor cortex despite a loss of RFP expression at 9 weeks post-PTEN-KO. However, few Fluorogold labeled neurons were detected within the motor cortex at 6 months post-treatment. BDA labeling from the motor cortex revealed a dense corticospinal tract (CST) bundle in all groups except chronically treated PTEN-KO mice indicating a potential long-term toxic effect of PTEN-KO to neurons in the motor cortex. PTEN-KO mice had significantly more β**-**tubulin III labeled axons within the lesion when treatment was delivered acutely, but not chronically post-SCI. In conclusion, we have found that using AAVrg’s to knockout PTEN is an effective manipulation capable of restoring motor functions in chronic SCI and can enhance axon growth of currently unidentified axon populations when delivered acutely after injury. However, the long-term consequences of PTEN-KO may exert neurotoxic effects.

## 1.0 Introduction

### 1.1 Preface

The intent of basic and pre-clinical scientific literature is designed to advance scientific knowledge and prospective treatment strategies. However, interactions between scientists and those living with spinal cord injuries have revealed that scientific literature is actively consumed by those with lived experience. It is therefore essential for the dissemination of scientific literature to be obtainable and understandable to both scientists and a lay audience that possess vested interests. In an active effort to appropriately engage a broader community, we have partnered with individuals with lived experience to formulate a layman’s summary of the project aims and key findings, which is provided at the end of the discussion section (section 3.1).

### 1.2 Introduction

Damaged axons do not regenerate within the injured central nervous system (CNS). The extent of spared axons after a spinal cord injury (SCI) limits functional abilities. Enhancing communication to-and-from the brain through induction of regeneration, axonal sprouting, or increased circuit excitability are all methods capable of restoring function after SCI and may be essential processes to restore function in chronic SCI (1, 2). Indeed, where-as use of acute neuroprotectants can improve outcomes when applied early after SCI, use of neuroprotectants in chronic injury conditions hold less merit as therapeutic approaches. Gainful manipulations to axons and neural circuitry may be required to improve neurological function in chronic SCI conditions.

Enhancing signaling through the PI3K/AKT/mTOR pathway is recognized as a mechanism capable of inducing axon growth and regeneration (3–7). Deletion of the phosphatase and tensin homolog protein (PTEN), which antagonizes PI3K activity, is sufficient to induce limited regeneration in corticospinal-(CST) and rubrospinal-tract axons after SCI (8–15). Inhibiting PTEN activity is identified actionable target capable of inducing axon growth when applied in both acute and chronic conditions of SCI (9, 14). Whether deleting PTEN from other spinal-projecting neurons can improve function after SCI remains unknown.

To date, most genetic manipulations to PTEN signaling have occurred via direct injection of adeno-associated viruses (AAVs) into the motor cortex or red nucleus to target single spinal tracts (e.g. the corticospinal or rubrospinal tracts) (9–11, 14). While targeting defined axonal populations is sufficient to study the effects on regeneration, the effects are limited to only a single spinal tract and may not be sufficient to elicit locomotor improvements. Use of AAVs that are pseudotyped for retrograde transport (AAVrg) can be injected at the lesion site and taken up by near-all neurons projecting to or below the level of the lesion, with a few notable exceptions including neurons of the caudal Raphe nucleus and Locus Coeruleus (16–18). The use of AAVs for retrograde targeting of spinal axons was first performed by Klaw and colleagues (2013) prior to the development of a designer AAVrg envelope protein capable of more efficient retrograde transport (17, 19). The advent of the AAVrg pseudotype has gathered much interest as a prospective gene-delivery method for SCI due to its ability to affect both damaged and spared fibers of most spinal-projecting neurons (18). To the best of our knowledge, most AAVrg applications have only targeted acutely injured axons. Whether chronically injured axons can be targeted to facilitate functional improvements is not well understood.

To determine the answers to these outstanding questions regarding the chronically injured spinal cord, we applied AAVrg to delete PTEN acutely and chronically after a complete T8 crush SCI. Through a series of AAvrg and tract-tracing experiments, we demonstrate that a single spinal injection of AAVrg’s can target neuronal populations that are otherwise challenging to reach via conventional stereotactic methods and that a single AAVrg injection is sufficient to restore motor functions when applied in both acute and chronic SCI. Despite observing functional improvements in both conditions, detectable effects on axon growth were only observed when applied immediately post-injury. We further identify that the human Synapsin1 promoter (hSyn1) is potentially affected by PTEN deletion suggesting a role of sustained growth in regulating synaptic proteins; coincidently we observed that long-term PTEN deletion may also exert progressive neurotoxic effects on neurons in the motor cortex. Together, our findings indicate that chronically injured axons can be targeted with AAVrg to facilitate functional recovery after severe SCI. Additional work is needed to determine whether AAVrg delivery can facilitate sustained functional recovery without neurotoxicity in chronic SCI.

## 2.0 Results

### 2.1 Experimental Design

To study the effects of PTEN-KO on axon growth and regeneration we utilized a complete crush model of SCI in PTEN^FloxΔ^/^Δ^ mice as described in prior literature (8, 9, 14, 20). Complete crush models leave limited spared axons through the lesion and are hallmarked by astrocyte infiltration into the lesion in chronic SCI, which is essential to support axon regeneration (14, 21). To target spinal-projecting neurons throughout the neuroaxis we delivered an AAvrg carrying cre recombinase and/or a red fluorescent protein (RFP), which were produced under the hSyn1 promoter.

AAVrg’s were intraspinally injected rostral to the lesion either immediately after SCI, or 3 months post-SCI. After SCI, locomotor abilities were analyzed every other week using the Basso Beattie and Bresnahan scale of locomotor recovery (BBB)(22). The BBB is typically used to assess locomotor recovery in rats; however, the more common Basso Mouse Scale (23) has a limited sensitivity at the lower ranges of functions, e.g. for animals without weight support which is anticipated after complete spinal crush (see methods section 5.3). Functional outcomes were obtained in mice treated either acutely or chronically after SCI. A separate group of acutely treated mice was used for both anterograde tracing of the CST using biotinylated dextran amines (BDA) or retrograde tracing of spinal-projecting neurons using Fluorogold. See Fig. 1 for a graphical summary of the experimental design. Groups, sample sizes, attrition rates, and outcomes are available in Table 1.

**Figure 1.**
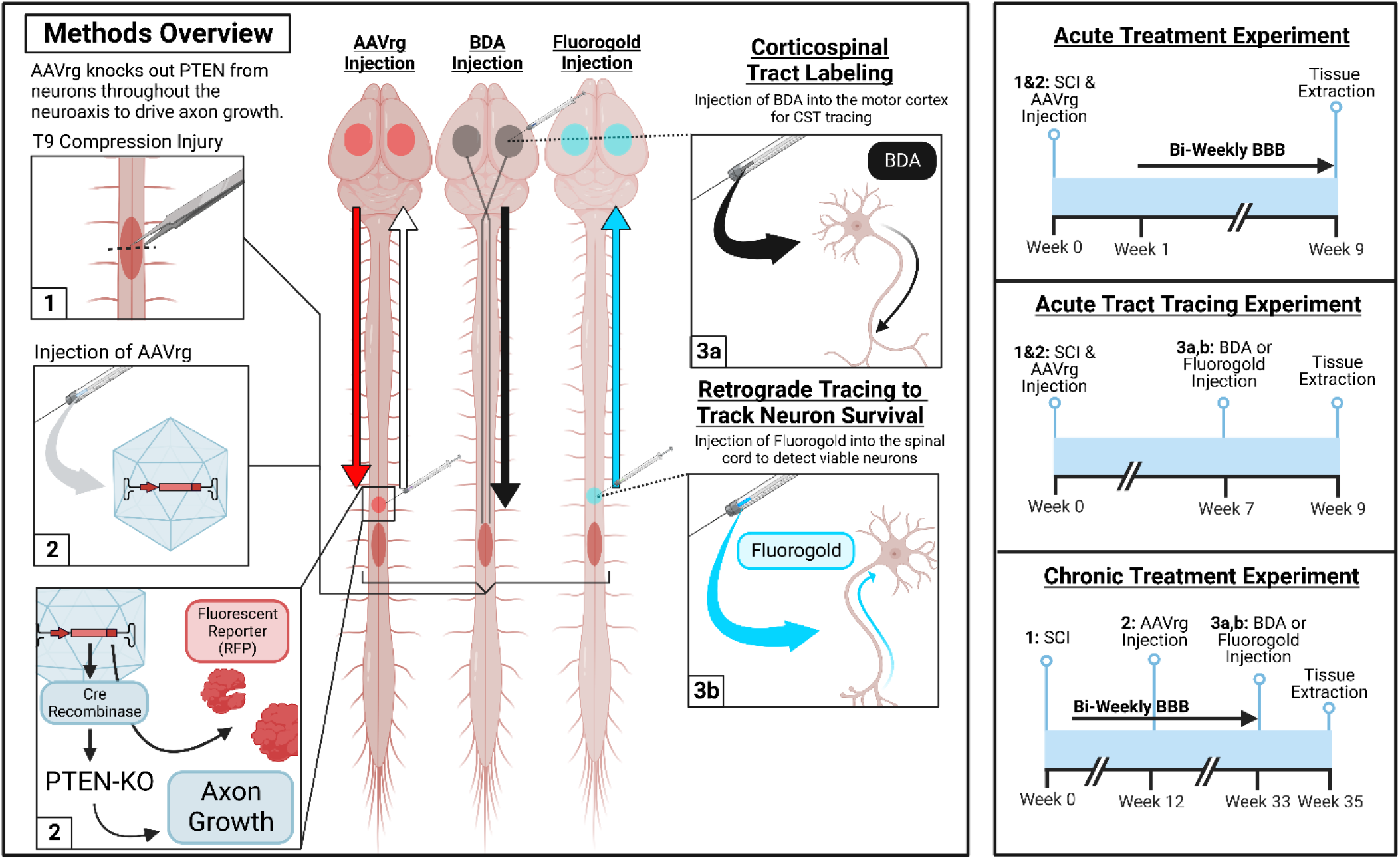
Overview of methods, experimental timelines, experimental groups, outcomes, and sample sizes. Three separate tracing procedures were performed throughout our experiments. First, AAVrg injections into the spinal cord retrogradely transport from the injection site into neurons throughout the brain. RFP reporter proteins are then anterogradely transported to fill axons above the lesion. Next, in a subset of mice, the corticospinal tract (CST) was traced by injecting biotinylated dextran amines (BDA; 10,000 kD) into the motor cortex to anterogradely label axons within the spinal cord. Finally, in another subset of mice, Fluorogold was injected into the spinal cord as a retrograde dye to detect surviving neurons throughout the brain, enabling visualization of neurons that lost RFP labeling. We utilized three major experiments to assess the efficacy of AAVrg’s to knockout PTEN as a treatment for SCI. First, male mice were treated at the time of SCI and were evaluated bi-weekly for behavioral analyses for up to 9-weeks post-injury. Next, both male and female mice were treated at the time of injury and were labeled with either BDA or Fluorogold at 7-weeks post-injury. Finally, female mice were treated with AAVrg’s at 12-weeks post-SCI, monitored for locomotor improvements bi-weekly, and either received BDA or Fluorogold labeling 9-months post-SCI. This image was created with BioRender.com.

**Table 1.**
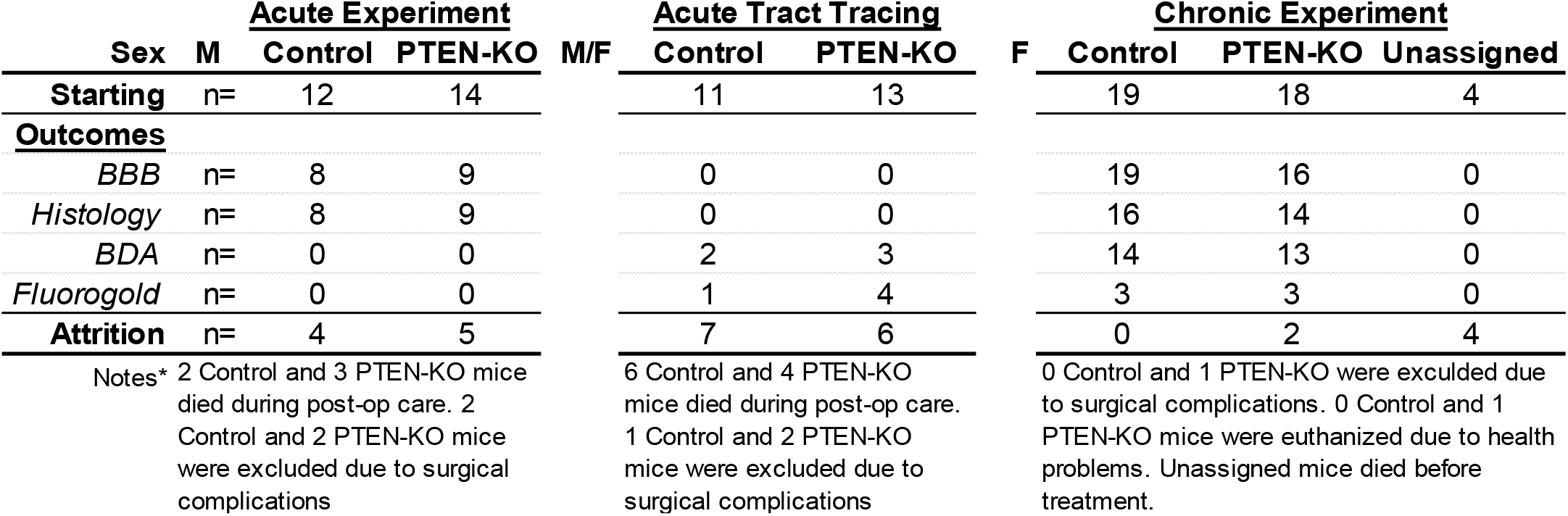
Experimental groups, outcomes, and sample sizes. Starting group sizes, final sample size counts, and attrition are reported for each outcome in each experiment. High attrition was observed in acute experiments which were caused in large part by bladder rupture within a few days post-SCI during daily bladder care in male mice.

### 2.2 Distribution and Fluorescent Expression of AAVrg’s Reveal a Consistent and Stable Labeling Throughout the Neuroaxis in Control but not PTEN-KO Mice

#### 2.2.1 AAVrg transduces neurons throughout the neuroaxis but with low efficacy in the Locus Coeruleus and caudal Raphe nucleus

Recently published reports which utilized spinal cord injections of AAVrg’s in the absence of injury or acutely after SCI have characterized the distribution of viral expression throughout the neuroaxis (16, 18, 24). Our experiment, however, specifically evaluated the expression patterns in neurons as long as 9 months post-SCI in chronic-treated conditions, utilized the hSyn1 promoter to limit expression to neurons, and evaluated the effects of sustaining PTEN-KO for up to 6 months.

When evaluating the labeling patterns in the brain we focused on a few main areas of interest identified through prior published reports (16, 18). Specifically, neuron labeling was evaluated in the motor cortex, red nucleus, Locus Coeruleus, Barrington’s Nucleus, as well as the caudal Raphe nucleus at 2 weeks post-injection in acutely injured spinal cords using AAVrg-hSyn1-eGFP. We validated that labeling could be detected at 2 weeks post-injection throughout the motor cortex, red nucleus, and Barrington’s nucleus (Fig. 2). Next, we validated that similar labeling patterns were observed at both 9 weeks and 9 months post-SCI in control AAVrg-treated mice (Figs. 3,4). At all time-points, co-labeling between DBH and eGFP within the Locus Coeruleus was scarce. We also observed scarce co-labeling of eGFP^+^ neurons with 5-HT within the Raphe nucleus cluster. Our results confirm previous reports demonstrating that AAVrg targets neurons throughout the neural axis with some limitations. Here we confirm these effects take place after SCI and are sustained for up to 9 months after injection (16, 18, 24).

**Figure 2.**
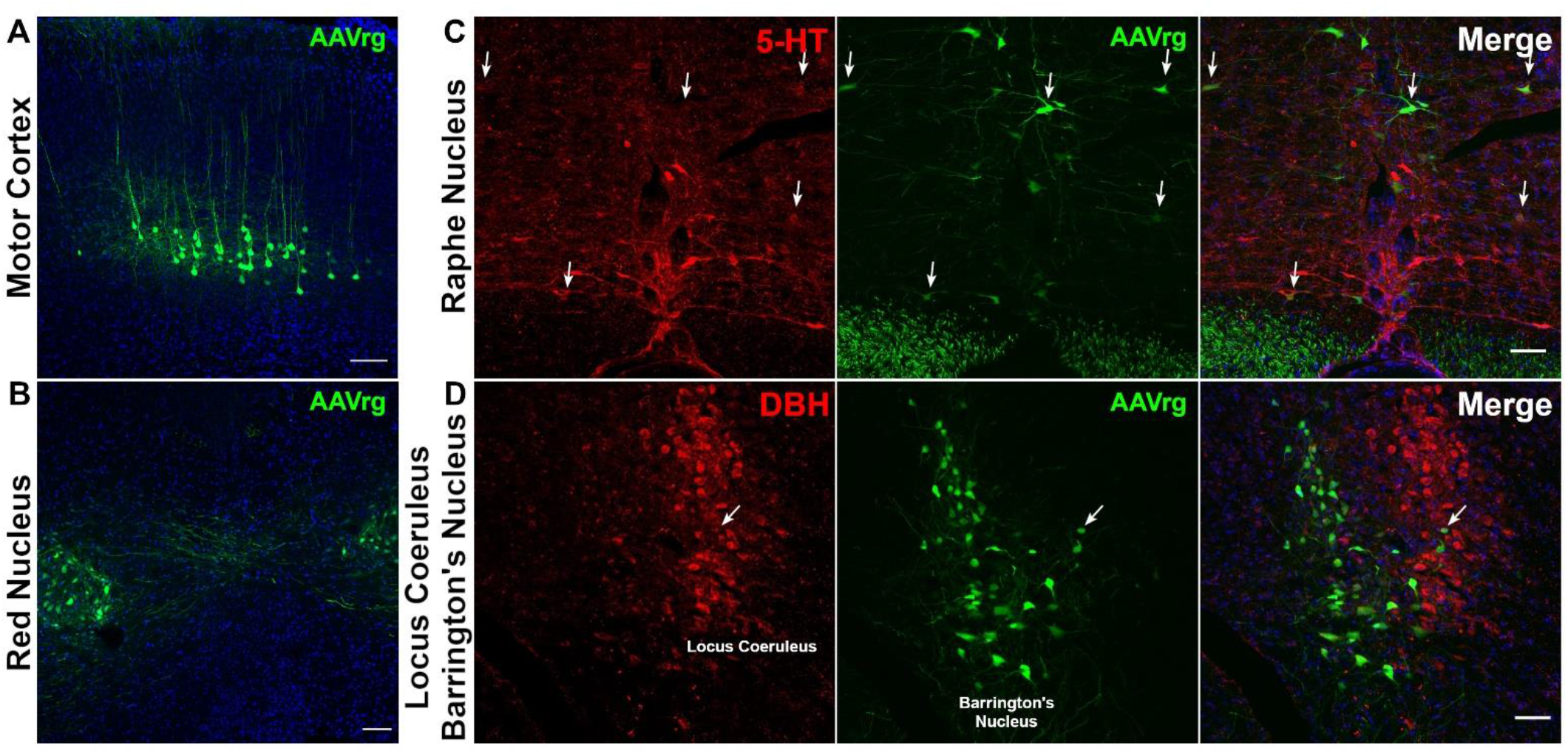
AAVrg injections into the spinal cord transduces neurons throughout the neuroaxis. A single injection of AAVrg-eGFP into the T8/9 spinal cord of acutely injured mice retrogradely transports to the cell bodies throughout the neuroaxis. At 2-weeks post-injection, neurons within the motor cortex (A), red nucleus (B), and Barrington’s nucleus (D) are observed to be transduced by the virus. In contrast, despite observing many infected neurons within the reticular formation only a few infected neurons within the caudal Raphe nucleus (C) are co-labeled with serotonin (5-HT). Further, very few infected neurons were co-labeled with dopamine beta hydroxylase (DBH). Arrows point to co-labeling between AAVrg and 5-HT or DBH. Blue labeling is Dapi. Scale bars = 100 μm.

**Figure 3.**
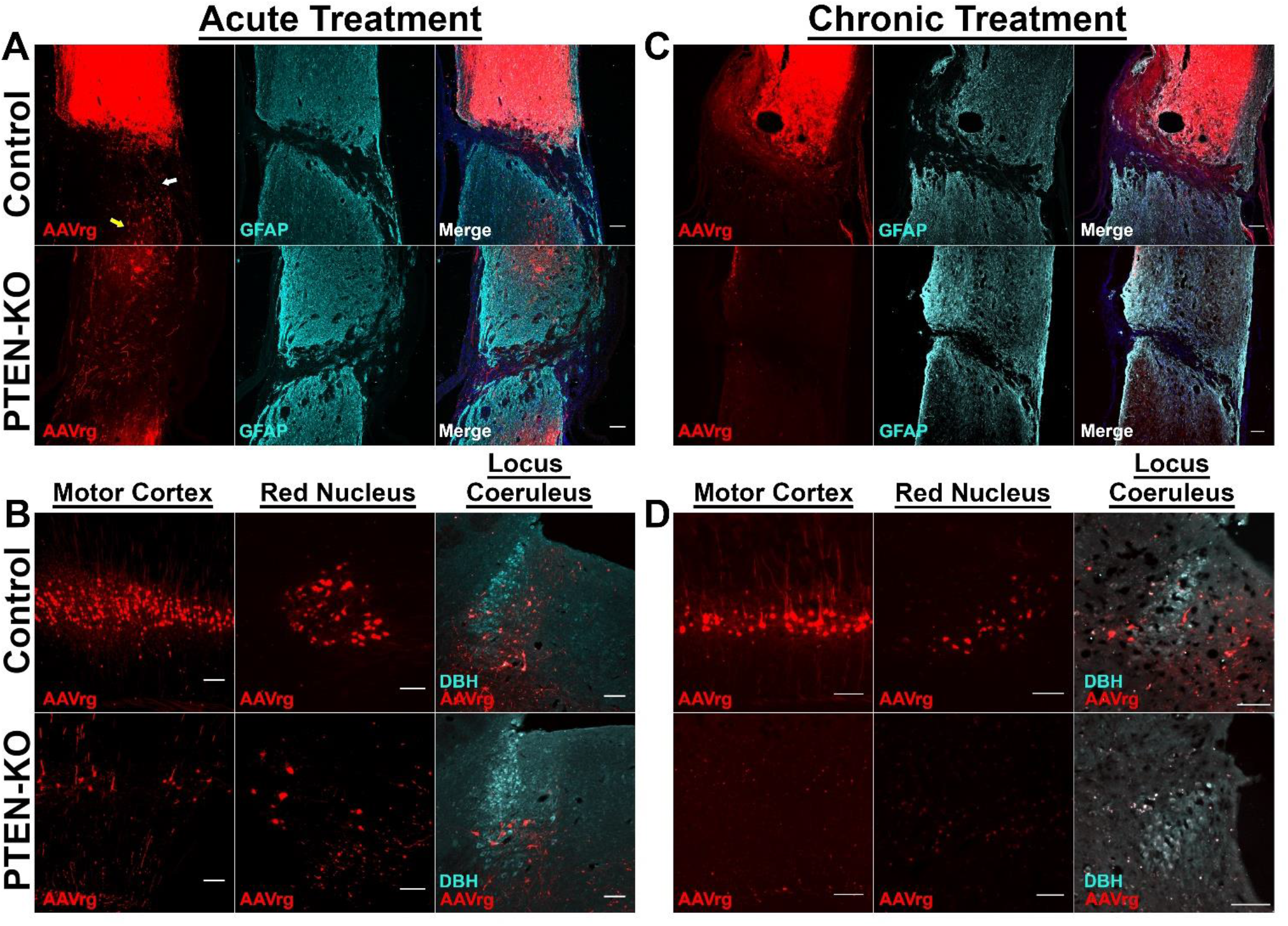
The RFP reporter that is expressed with the hSyn1 promoter is diminished after PTEN-KO. SCI mice were treated with AAVrg that expressed an RFP alone, or with Cre-recombinase to knockout PTEN under the hSyn1 promoter, and survived for 9-weeks (treated at the time of SCI) (A,B), or 6-months post-injection (treated at 12-weeks post SCI)(C,D). At both 9-weeks and 6-months post-injection RFP expression remained stable and was detected in all of the same regions as displayed in Fig. 2 in control-treated mice. In contrast, RFP expression was diminished in the spinal cord (A), and brains (B) by 9-weeks post-injection in PTEN-KO, and was almost completely absent by 6-months post-injection (B,D). White and yellow arrows (A) point to evidence of spared axons across the lesion and transduced neurons caudal to the lesion respectively. Both the existence of spared fibers and transduced neurons limit the ability to use RFP caudal to the lesion as a metric of axon regeneration. Blue labeling is Dapi. Dopamine beta hydroxylase (DBH). Scale bars = 100 μm.

**Figure 4.**
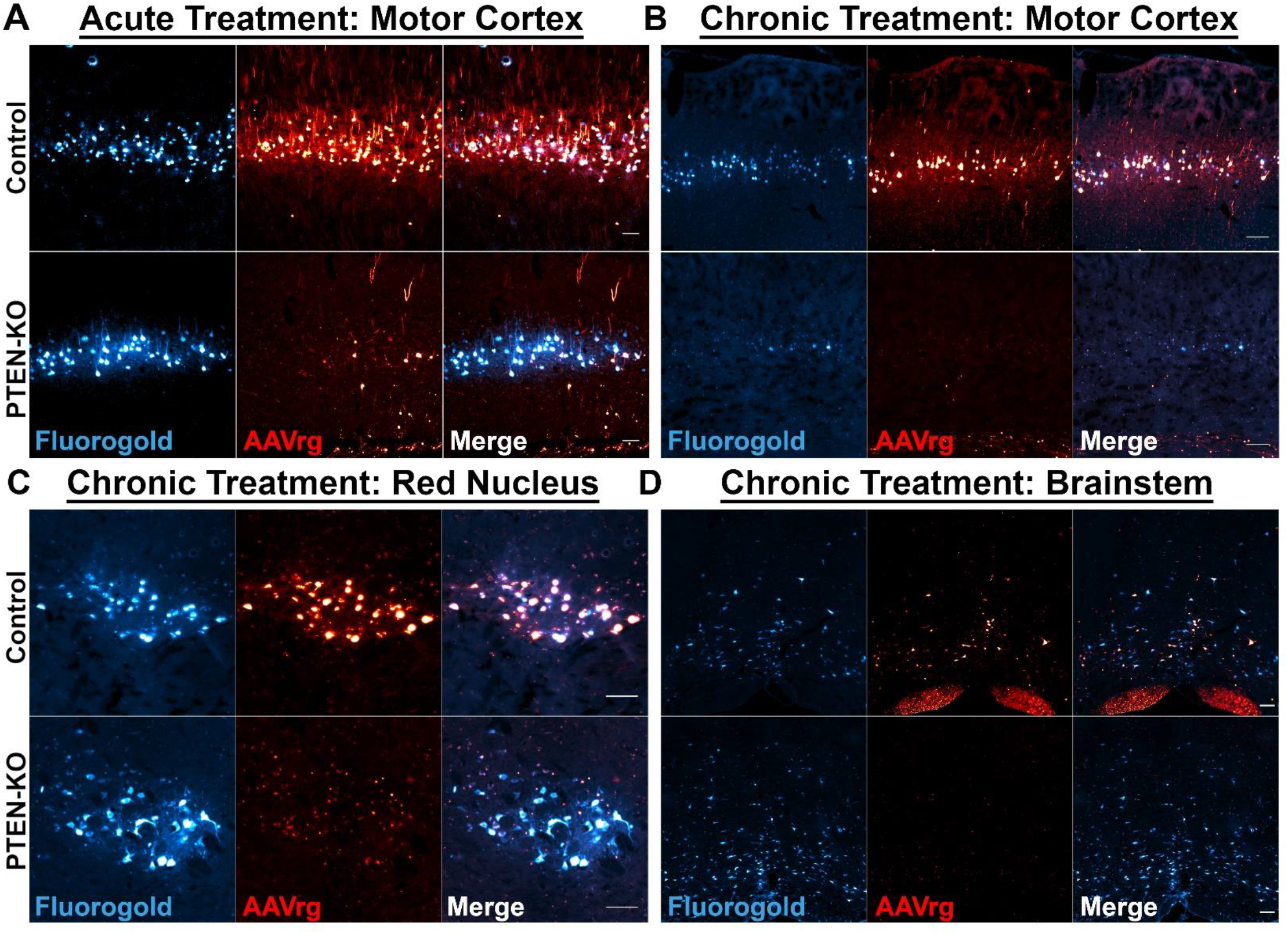
Co-labeling with Fluorogold and AAVrg-RFP reveals a PTEN-KO induced loss of neurons in the motor cortex at 6-months, but not 9-weeks post-treatment. A unique loss in RFP expression in PTEN-KO mice created a question regarding the viability of transduced neurons. To address questions about viability, the spinal cords of mice were injected with Fluorogold rostral to the lesion, 2-weeks prior to euthanasia. The motor cortex was evaluated at 9-weeks post-SCI following acute treatment and revealed a preservation of neuron densities, suggesting that neurons remain viable after PTEN-KO and that mechanisms other than neuronal loss underlie the loss of RFP expression (A). However, by 6-months post-PTEN-KO in mice treated at 12 weeks after SCI there was an apparent loss of neurons labeled with Fluorogold in the motor cortex relative to controls (B). Fluorogold was still observed within the red nucleus (C) and throughout the brainstem (D) at 6-months post-treatment. A loss of Fluorogold in the motor cortex at 6-months, but not 9-weeks post-treatment, points to a potential toxic effect of sustained PTEN-KO in specific neuronal populations including the motor cortex. Note: Most of the red puncta in the chronically treated PTEN-KO images (B,C,D) is caused by an accumulation of autofluorescence (i.e. lipofuscin), rather than viral expression. Some RFP expression is detected in the corpus callosum in chronically treated tissue (B). Scale bars = 100 μm.

#### 2.2.2 Reporter gene expression under the hSyn1 promoter is diminished in neurons after PTEN-KO

Next, we compared labeling of patterns between control- and PTEN-KO-treated mice within the brains and spinal cords. After SCI, at both analyzed time points, reporter gene labeling in control-treated mice saturated the injection site making it difficult to capture and image finely labeled fibers within the injury epicenter. Labeling was found in axons within the white matter and neuronal cell bodies within the gray matter rostral to the lesion. Further, we identified axons spanning the lesion site primarily in the ventral aspect of the spinal cord. These axons may indicate that although the crush lesions were severe, many were not anatomically complete, or are indicative of limited sprouting/regeneration after injury (White arrow; Fig. 3A).

Axons that spanned the lesion were of similar morphology to what would be expected of axon sparing, rather than regeneration (25). Further, we observed the existence of cell bodies caudal to the lesion which could have been labeled either due to having axons that were spared from injury, potential leakage of the virus across the lesion site, or could represent axons that took up the virus immediately after injury and transported the viral particles to the soma prior to becoming axotomized during secondary injury cascades (Yellow arrow; Fig. 3A). Due to evidence of spared axons and labeled cell bodies caudal to the lesion, we are unable to use reporter gene expression below the injury as a measure of axon regeneration and will refer to the lesion severity as severe and near-complete.

In mice which received PTEN-KO, fluorescent labeling was drastically reduced. Scant neurons were labeled throughout the analyzed locations by 9-weeks post-treatment and labeled neurons appeared hypertrophic (Fig. 3B) as previously reported with PTEN inhibition (8, 14, 20). By 9-months post-PTEN-KO, AAVrg-mediated fluorescent was extremely limited or non-existent in every location (Fig. 3).

#### 2.2.3 PTEN-KO exerts signs of long-term toxicity to neurons in the motor cortex at 6-months, but not 9-weeks post-injection

To address if PTEN-KO was leading to neuronal death, we performed retrograde labeling in a subset of SCI animals using Fluorogold at 9 weeks post-injection. In both PTEN-KO and control mice treated acutely post-SCI, we detected Fluorogold labeling of neurons throughout the neural axis in locations such as the motor cortex (Fig. 4A), suggesting that neuronal viability remained at 9-weeks post-treatment. To determine the long-term effect of PTEN-KO on neuron viability we performed track tracing 6 months after injection following treatment in mice with chronic SCI (3 months post-injury). In contrast to 9-weeks after injection, at 6 months post-AAVrg injection (9 months post-SCI) in PTEN-KO mice, we failed to detect strong Fluorogold labeling in the motor cortex, potentially indicative of reduced neuronal viability with sustained PTEN-KO (see below) (Fig. 4B). Interestingly, however, we observed a conserved density of Fluorogold-labeled neurons in the red nucleus and throughout the brainstem in PTEN-KO mice at this 6 month timepoint (Fig. 4C,D).

Taken together we have validated that the use of AAVrg’s does indeed expand the gene-therapy effect to a wider range of neurons throughout the neuroaxis. However, we have identified two confounding effects when performing PTEN-KO: 1) PTEN-KO may silence the hSyn1 promoter which is both consistent with a need to reduce synaptic strength for regeneration (26–30); and 2) sustaining long-term growth via PTEN-KO may have adverse effects on neuronal health and viability of specific populations of neurons.

### 2.3 Improvements in Hind-Limb Function After PTEN-KO are Observed in Both Acute and Chronic SCI

#### 2.3.2 Acute PTEN-KO using AAVrg’s restores weight supported stepping

To determine if PTEN-KO using AAVrg’s could improve functional recovery, we first applied AAVrg’s above the lesion immediately post-SCI and monitored for locomotor recovery bi-weekly for 9-weeks. Relative to control-treated mice, PTEN-KO significantly improved hind-limb function with most mice displaying a robust increase in range of motion, sweeping behaviors, or plantar placing (treatment effect, F(_1,15_)=16.47, *p*=0.001; time by treatment interaction, F(_4,60_)=9.107, *p*<0.0001)(Fig. 5A). Several mice regained weight supporting abilities sufficient to perform at least occasional overground stepping (Fig. 5B). In contrast, control-treated mice had no recovery of hind limb function and displayed only minimal range of motion usually within the ankle joint. When characterizing the behavioral return there was an apparent increase in function beginning after 5-weeks post-treatment (Fig. 5B).

**Figure 5.**
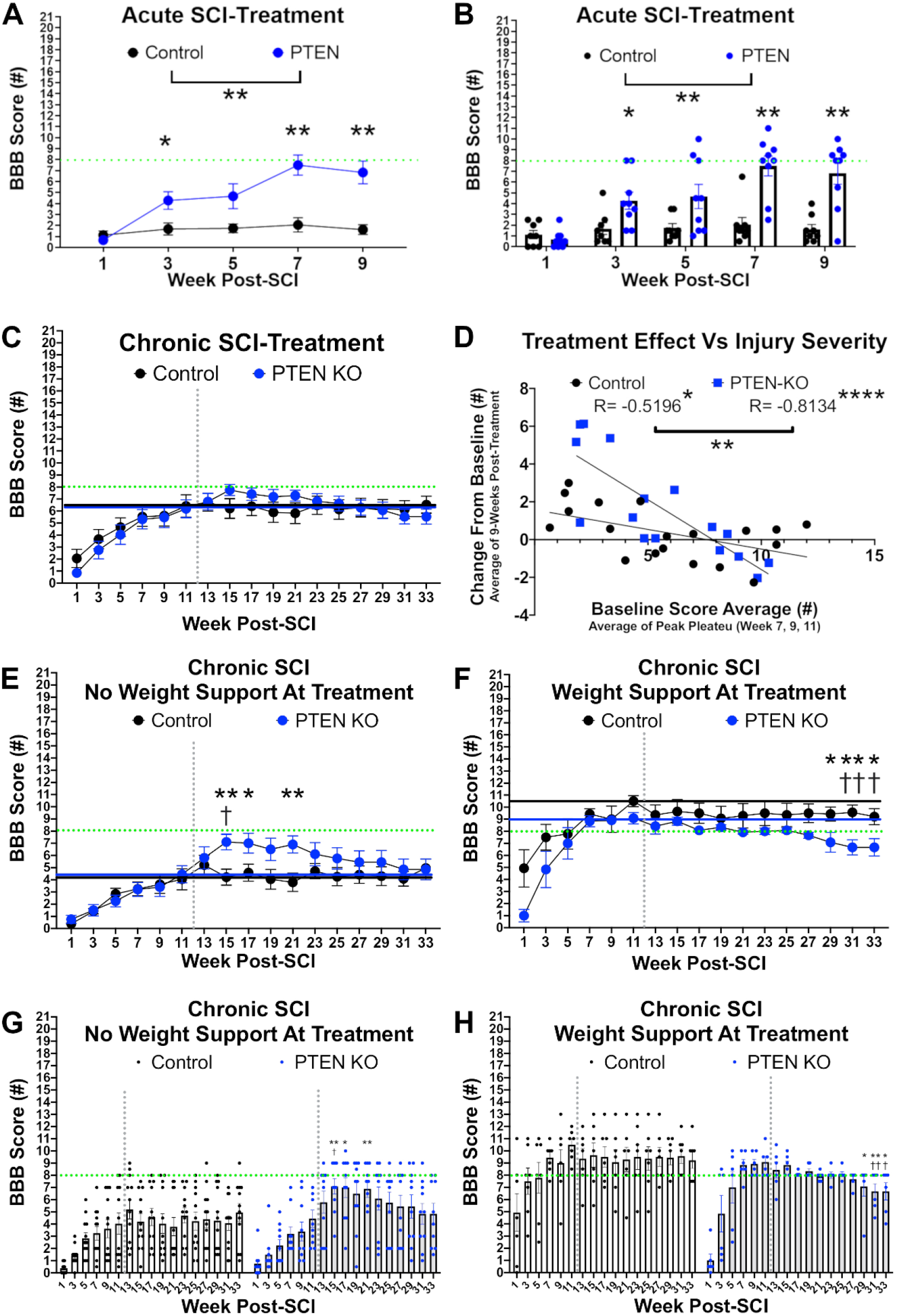
PTEN-KO using AAVrg improves locomotor outcomes when delivered at both acute and chronic timepoints but not in mice with pre-existing weight support. Mice were treated with either a control AAVrg-RFP, or AAVrg-Cre-RFP to knockout PTEN either immediately post-SCI (A,B) or at 12-weeks post-SCI (C-H). PTEN-KO significantly improved hind limb abilities in mice treated acutely post-SCI with several mice regaining weight supporting abilities (F_(1,15)_=16.47, *p*=0.001; A,B). PTEN-KO exerted a heterogeneous effect when delivered in chronic SCI. While a significant time-by-treatment interaction was observed (F_(16,528)_=1.845, *p*=0.023; C), no individual time point was significant using pair-wise comparisons when looking at all PTEN-KO-treated mice together. However, there was a strong significant negative correlation between the treatment effect and the functional abilities at time of treatment (control mice, R=-0.5196, *p*=0.022; PTEN-KO mice, R=-0.8134, *p*=0.0001; D) with the slopes of the regression lines differing significantly by group (F_(1,31)_=11.55, *p*=0.0019). Several of the most severely injured mice regained weight supporting abilities for the first 9-weeks post-treatment (E), where-as many mice with weight supporting abilities at the time of treatment lost the ability to weight support (F). To account for pre-treatment abilities being a significant co-variate that introduced a divergence in our analyses, for treatment initiated 12-weeks after SCI, we separated mice into pre- and post-weight supporting abilities at time of treatment. After separating mice into pre- and post-weight supporting groups, we observed a significant time-by-treatment interaction (F_(16,176)_=1.845, *p*=0.0287) in pre-weight supported groups. PTEN-KO mice exhibited both within and between group improvements within the first 9-weeks post-treatment relative to baseline and control treated mice respectively. In contrast, most mice with weight support at the time of treatment lost weight support within the first 9-weeks post-treatment. Regardless of how mice were split for analysis, after 9-weeks post-PTEN-KO, mice began to lose function progressively until the end of the study at 33-weeks post-SCI. Green dotted horizontal line = the boundary between weight support and no weight support on the BBB scale (A-C, E-H). Grey dotted vertical line = time of treatment (C, E-H). Repeated-measures ANOVA was performed using either Fisher’s LSD for between-group post hoc comparisons or Dunnett’s pair-wise comparisons for comparing within group scores to pre-treatment values (A-C, E-H). Graphs B,G,H are provided to display individual group distributions. Sample sizes; control = 8, PTEN-KO = 9 (A-B); control = 19, PTEN-KO = 16 (C-D); control = 12, PTEN-KO = 10 (E,G); control = 7, PTEN-KO = 6 (F,H). Error bars are S.E.M. Between groups comparisons = \**p* < 0.05, \*\**p* < 0.01, \*\*\**p* < 0.001, \*\*\*\**p* < 0.0001. Within group comparisons for PTEN-KO mice = ^†^*p* < 0.05, ^††^*p* < 0.01.

#### 2.3.3 Chronic PTEN-KO using AAVrg’s exerts injury-severity dependent effects

To determine whether PTEN inhibition could improve function after chronic SCI, we delivered control or PTEN-KO AAVrg in mice 3 months after SCI. Animals were randomly assigned to balanced treatment groups based upon locomotor function prior to treatment. We then evaluated locomotor performance bi-weekly for up to 6-months post-treatment (Fig. 5C-H).

In contrast to the acutely treated mice, chronically treated mice displayed complicated behavioral changes in response to PTEN-KO. At first glance of the data when all mice are analyzed in their respective groups, we do not see a main effect of PTEN-KO on motor performance but did detect a significant time-by-treatment interaction (treatment effect, F(_1,33_)=0.006, *p*=0.937; time by treatment interaction, F(_16,528_)=1.845, *p*=0.023)(Fig. 5C). Upon further analyses, we noticed that mice with weight-supporting abilities (high function) at the time of PTEN-KO lost weight-supporting abilities after treatment. In contrast, mice with severe and near-complete paralysis (low function) at the time of PTEN-KO gained significant locomotor functions with several mice regaining weight support (Fig. 5).

We performed a linear regression to see if the locomotor functions at the time of AAVrg injection could predict a treatment response and observed a significant negative correlation in the PTEN-KO mice between the BBB score before treatment and the change in BBB score averaged over the first 9-weeks after treatment (control mice, R=-0.5196, *p*=0.022; PTEN-KO mice, R=-0.8134, *p*=0.0001). To ensure the effects we were observing were due primarily to the PTEN-KO as opposed to the surgical injection of AAVrg’s, we compared the regression lines between control and PTEN-KO groups and found the slopes to be significantly different (F(_1,31_)=11.55, *p*=0.0019)(Fig. 5D), further validating that the PTEN-KO mice experience a severity-dependent treatment effect. Most importantly, we observed the regression line crossing the X axis around a BBB score of 8, which marks the transition from no to some-hindlimb weight supporting abilities. In other words, the results from our regression analyses indicate that hindlimb weight-supporting abilities are disrupted after PTEN-KO in high-functioning injured mice, but reciprocally, lower-functioning mice experienced a therapeutic benefit.

To further investigate these severity-specific treatment effects, we analyzed high (>8) and low (<8) functioning mice separately. Relative to high-function mice that received control AAVrg, high function mice with PTEN-KO had a significant decrease in performance after treatment with a gradual exacerbation of deficits over time that resulted in a significant time-by-treatment interaction (F(_16,176_)=1.845, *p*=0.0287; Fig. 5F). All high function mice treated with PTEN-KO using AAVrg’s lost weight supported stepping abilities by the end of the study (Fig. 5H). In contrast, locomotor function significantly improved in low-function mice receiving PTEN-KO relative to low-functioning controls, indicated by a significant time-by-treatment (F_(16,320)_=1.715, *p*=0.0426; Fig. 5E). Several low functioning PTEN-KO mice gained weight supported stepping abilities (Fig. 5D,G). Importantly, the functional gains of low-function animals with PTEN-KO began to return to pre-treatment levels 9-weeks post-treatment with all gains lost within 6 months of treatment (Fig. 5D).

### 2.4 PTEN-KO Exerts Different Effects on Axon Growth Between Acute and Chronic SCI

#### 2.4.1 PTEN-KO augments axon growth into the lesion in acute but not chronic SCI

Our original intent was to characterize axon growth and axon regeneration by determining RFP labeling within and beyond the lesion. As mentioned above this was compromised by a PTEN-KO-specific loss of RFP signal. While we have quantified RFP labeling within the lesion, results should be interpreted according to these major limitations. Instead of relying on viral RFP expression, we labeled sections against β-Tubulin III (β3-Tub) to visualize all axons within the lesion. The lesion boundaries were traced, and the proportional area covered in β3-Tub labeling was assessed as a total axon growth marker. At 9-weeks post-SCI in acutely treated mice, β3-Tub^+^ area was significantly increased within the lesions of PTEN-KO mice, displaying a 2-fold increase relative to controls (control M=14.07, PTEN-KO M=30.27; T_(16)_=3.141, *p*=0.0067)(Fig. 6A,C). In contrast, 6 months after treatment in chronically injured mice, β3-Tub^+^ area was not significantly increased in the lesions of PTEN-KO mice relative to controls (control M=10.37, PTEN-KO M=12.67; T_(28)_=1.037, *p*=0.308)(Fig. 6B,D).

**Figure 6.**
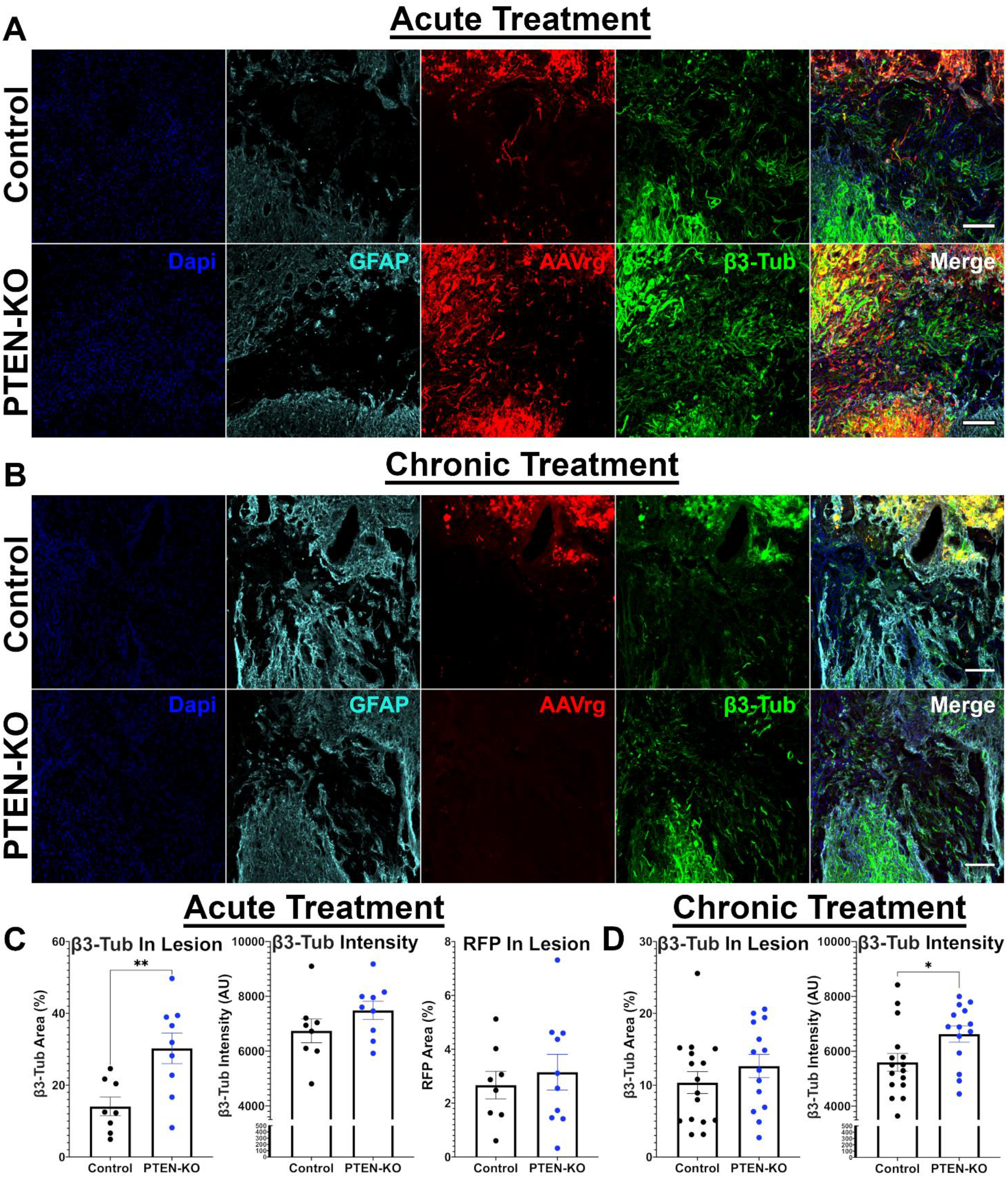
Using AAVrg to knockout PTEN increases β3-Tub axon labeling within the lesions of acutely, but not chronically, treated mice. Spinal cords were extracted from mice at either 9-weeks or 9-months post-SCI that were treated with either a control AAVrg-RFP, or AAVrg-Cre-RFP. Spinal cords were extracted at 9-weeks post-SCI in mice treated acutely post-injury, or at 6-months post-treatment in mice treated at 12-weeks post-SCI. Spinal cords were labeled for GFAP to identify the glial scar, as well as β-tubulin III (β3-Tub) to identify axons within the lesion (A,B). As previously described, RFP expression was diminished in PTEN-KO mice at 9-weeks post-treatment and was almost completely absent at 9-months post-treatment. The percentage β3-Tub labeling within the lesion was quantified as a measure of axon growth. Further, the labeling intensity of positive β3-Tub pixels was determined. A similar analysis was performed for RFP fluorescence in tissue obtained from acutely treated mice but was not performed for tissue obtained from chronically treated mice due to an absence of fluorescence. Lesions in the spinal cords of acutely treated mice presented with a significant increase in proportional β3-Tub area (T_(16)_=3.141, *p*=0.0067), but no differences were observed for the intensity of β3-Tub labeling (T_(16)_=1.355, *p*=0.194) or the proportional area of RFP within the lesion (T_(16)_=0.55, *p*=0.589; C). Lesions in the spinal cords of chronically treated mice did not differ in the proportional area of β3-Tub labeling within the lesion (T_(28)_=1.037, *p*=0.308), however the intensity of β3-Tub labeling was significantly brighter (T_(28)_=2.308, *p*=0.028; D). Unpaired T-Tests were used for statistical comparisons. Scale bars = 100 μm. Error bars are S.E.M. Sample sizes; control = 8, PTEN-KO = 9 (C); control = 16, PTEN-KO = 14 (D). \**p* < 0.05, \*\**p* < 0.01.

Our observation of increased axon growth into the lesions of PTEN-KO mice at acute, but not chronic, time points are consistent with previous findings that assert treatment strategies aimed at inducing regeneration have less efficacy when delayed into chronic settings (9). However, while we did not observe a total increase in β3-Tub^+^ area in PTEN-KO mice in chronic SCI, we did observe a difference in the labeling intensity. The β3-Tub^+^ area within the lesion had a brighter and more vivid labeling intensity in PTEN-KO mice compared to controls in the chronic (T_(28)_=2.308, *p*=0.028)(Fig. 6D), but not acute experiment (T_(16)_=1.355, *p*=0.194)(Fig. 6C). This suggests that while total axon growth might not have been enhanced from PTEN-KO in chronic SCI, there are potential neurobiological differences occurring within the axons. For RFP labeling within the lesion, no differences were found at 9-weeks following acute treatment (T_(16)_=0.55, *p*=0.589)(Fig. 6C); labeling was not quantified at 9-months after chronic treatment due to an almost complete absence of RFP expression in PTEN-KO mice (Fig. 6D).

#### 2.4.2 No differences in 5-HT axons are found across the lesion in PTEN-KO mice at either time point

Next, we evaluated the regenerative effects on 5-HT axons. While there was growth of 5-HT axons into and beyond the lesion even in control mice, there was no difference in 5-HT axon labeling at 9-weeks post-treatment following acute treatment (main effect, F_(1,10)_=0.004, *p*=0.948). Further, while 5-HT axon growth was observed in several animals from both groups, not every animal displayed such effects. Specifically, at 9-weeks post-SCI 5 of 8 control-treated, and 7 of 9 PTEN-KO treated mice had evidence of 5-HT growth caudal to the lesion (χ^2^=0.476, *p*=0.49), and this growth was limited to 400 µm caudal to the lesion border at the longest observed fiber (Fig. 7A,C).

**Figure 7.**
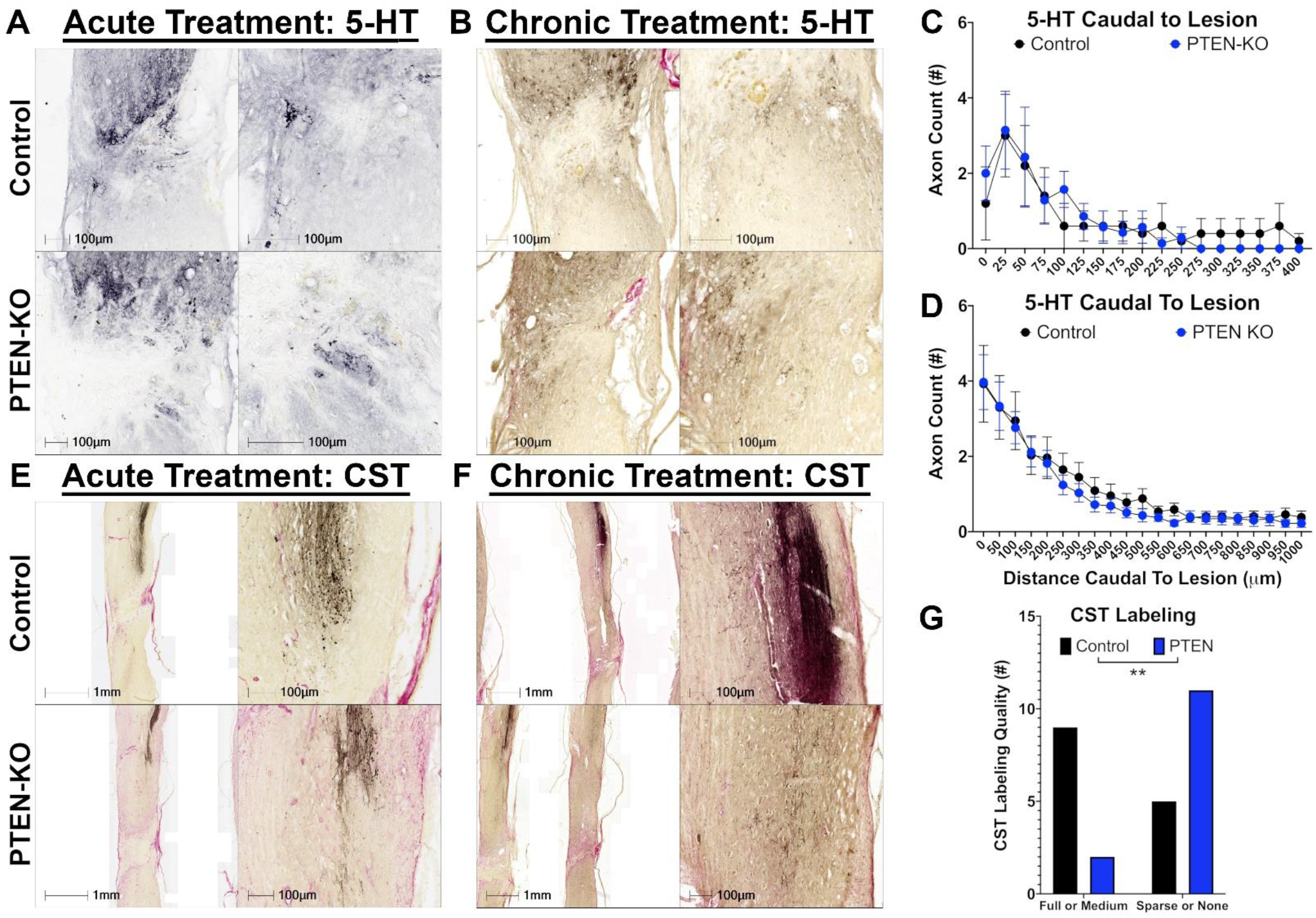
PTEN-KO using AAVrg did not regenerate serotonergic axons or CST axons when treated either acutely or chronically post-SCI. Immunohistochemistry was performed to visualize serotonergic (5-HT) axons to identify regeneration of supraspinal-projecting axons arising from the caudal Raphe nucleus (A). Further, anterograde labeling using biotinylated dextran amines (BDA) was performed from the motor cortex to label the corticospinal tract (CST; E,F). While 5-HT fibers were observed crossing the lesions in tissue obtained from both acutely and chronically treated mice at 9 weeks and 6 months after treatment respectively, there was no evidence that PTEN-KO enhanced axon regeneration at either time point (C,D). While the CST was only successfully labeled in 2 acutely treated control and 3 PTEN-KO mice using BDA, there was no evidence of axon growth or regeneration beyond the lesion in any analyzed tissue section (E). Surprisingly, in the 14 chronic PTEN-KO-treated mice that were labeled with BDA, there was an apparent loss of BDA labeling in most samples (F). BDA labeling was qualitatively assessed as either full labeling quality, medium labeling quality, sparse axon labeling, or no axon labeling. Ratings were grouped into either full or medium labeling, and sparse or no labeling for analysis, the results of which support a loss of CST labeling in PTEN-KO mice at 6-months post-treatment (χ^2^=6.677, *p*=0.0098)(G). A loss of neurons in the motor cortex (Fig. 4) that comprise the CST is a likely explanation for reduced BDA labeling in PTEN-KO mice. Scale bars are labeled as either 1 mm, or 100 μm. Error bars are S.E.M. Repeated-measures ANOVA was used for statistical analysis (C,D). Chi-Squared used for statistical analysis (G). Sample sizes; control = 5, PTEN-KO = 7 (C); control = 15, PTEN-KO = 13 (D); control = 11, PTEN-KO = 10 (G). \*\**p* < 0.01.

In contrast to our acute experiments, all chronically treated had evidence of 5-HT growth caudal to the lesion 9 months after SCI and 6 months after treatment. Importantly, some mice had evidence of spared 5-HT fibers which were detectable via 5-HT axon sprouting throughout the grey matter caudal to the injury in the lumbar cord (furthest caudal tissue evaluated) (Fig. 7B,D). However, similar to our acute experiment, there were no differences in axon regeneration between groups (main effect, F_(1,26)_=0.203, *p*=0.656; Fig. 7,D). Not observing significant differences in 5-HT axon regeneration between groups is not surprising considering the low number of 5-HT neurons successfully transduced with AAVrg in the caudal Raphe-Nucleus (Fig. 1C)(16, 18, 24).

#### 2.4.3 Corticospinal tract axons did not regenerate after PTEN-KO in either acute or chronic conditions

Due to a loss in RFP expression in PTEN-KO mice, as well as evidence of sparing and labeling of neuronal soma caudal to the lesion, we needed a method to evaluate regeneration specific to supra-spinal neurons. We performed BDA tracing from the motor cortex to determine the regenerative effects of PTEN-KO on the CST. Contrary to prior reports, we did not observe CST regeneration across the lesion in any of the animals at any time point, or in any group (Fig. 7E,F). Surprisingly, we did observe that most chronically treated PTEN-KO mice had poor or absent labeling of the CST (Fig. 7F).

To quantify our observation that PTEN-KO mice had less CST axons labeled with BDA at 6-months post-treatment, we rated CST labeling based on being visually completely filled, moderately filled, sparse evidence of axons, or no evidence of axons, and clustered animals into two groups for comparison using Chi-Squared. Mice with evidence of the CST being completely filled or moderately filled comprised one cluster, and mice with sparse evidence of axons or no evidence of axon labeling were clustered together. There were significantly less PTEN-KO treated mice with evidence of complete or moderate filling of the CST (χ^2^=6.677, *p*=0.0098)(Fig. 7G). Concerned that chronic PTEN-KO may be neurotoxic, we performed Fluorogold tracing in our last cohort of mice. As indicated in Fig. 4, the reduction in Fluorogold-labeled neurons in the motor cortex may explain the lower density and prevalence of CST labeling in mice with PTEN-KO at 6-months post-treatment (Fig. 4,7).

### 2.5 Long Term Health and Adverse Outcomes

Throughout both acute and chronic experiments we observed one unanticipated outcome that was specific to the PTEN-KO group. In mice receiving PTEN-KO in either the acute or chronic study, most-to-all mice receiving treatment developed an elevated chronic muscular tone of the abdominal muscles which we were not able to appropriately quantify, and therefore are only reporting through an empirical observation (for a photographic example see Supplementary Materials 1). An increased abdominal tone phenotype was apparent to the touch during routine bladder care. While we have not obtained quantitative evidence to support this claim, we believe that this hypercontraction of the abdomen was the cause of the higher functioning mice treated in the chronic condition to lose weight-supporting abilities after treatment. It is possible that mice maintaining a hyper-contracted abdomen experienced a disrupted ability to balance and maintain weight support. While we did see mice with the most severe lesions regain weight-supporting abilities in both the acute and chronic conditions, an increased abdominal tone could potentially support the ability of these severely injured mice to be capable of bringing their limbs under their body to assist in the recovery of weight supporting abilities. Importantly, when considering our AAVrg delivery approach, we injected the virus into the T8-9 vertebral segments which undoubtably affects neurons that control the abdomen. It is therefore important to consider that injection of viral vectors into the spinal cord does not just affect long-tract spinal projecting neurons but also affects neurons in the vicinity of the injection.

### 3.0 Discussion

There are several important and novel findings accumulated from work in this manuscript. 1) First, we have replicated that delivery of AAVrg’s can exert a wide genetic influence throughout the neuroaxis when applied rostral to an SCI lesion but that transduction efficacy within the caudal Raphe nucleus and Locus Coeruleus remain limited. 2) Even in severe and near-complete crush lesions, PTEN-KO using AAVrg’s improves locomotor outcomes in both acute and chronic SCI conditions, however, the effects are severity dependent. 3) PTEN-KO using AAVrg’s does increase axon growth into the lesions of acutely injured mice, demonstrating a growth effect on axons. 4) However, driving the RFP fluorescent reporter gene under the hSyn1 promoter revealed an unexpected interaction between the viral gene expression and PTEN-KO that diminished the RFP reporter. 5) Finally, we observed a potential toxic response to prolonged PTEN-KO in motor neurons comprising the CST, which is corroborated by a progressive loss in motor abilities over time, a loss of Fluorogold labeling within the motor cortex, as well as an absence of normal CST labeling within the spinal cords of PTEN-KO mice.

Use of gene delivery approaches to induce axon regeneration in SCI is an emerging strategy that has demonstrated preliminary success (30). To date, most gene-delivery approaches to manipulate PTEN have been performed with the intent of understanding the biological capacity for specific spinal tracts to regenerate, mainly the CST and rubrospinal tract. Use of retrogradely transported viral particles to target a larger breadth of spinal tracts via injection directly into the spinal cord confers major advantages to both study-specific spinal tracts of interest, but also to target neuronal populations that are otherwise difficult to affect. Prior work by both Blackmore and colleagues (16) and Metcalfe and colleagues (18) have characterized the capability for the AAVrg pseudotype to affect neurons and have found similar results, both concluding that glutamatergic neurons are very efficiently transduced, while neurons arising from the Raphe nucleus and Locus Coeruleus display limited transduction. We observed similar trends in the brains analyzed from our experiment, however, we did observe that a few labeled neurons in the Raphe nucleus were co-labeled with 5-HT. Additionally, we observed that use of the hSyn1 promoter is stable for up to 9-months post-transduction in every population of neurons analyzed, but the stability of this promoter may be dependent upon the regenerative state of the neuron.

PTEN-KO leads to increased activity of AKT and the mammalian target of rapamycin (mTOR) which exert modulatory roles across neuronal physiology. PTEN-KO leads to hyperexcitability of neurons and can induce epilepsy or seizures when occurring throughout the cortex (31). Effects of sustained hyper-activity of AKT/mTOR has also been found to induce spinogenesis on affected neurons and induce the formation of more synaptic contacts with pre-synaptic neurons (32, 33), as well as facilitate sprouting of axons to make more synaptic connections with post-synaptic neurons (8, 9, 11, 14). Taken together, both mechanisms are independent of long-distance axon regeneration but may exert therapeutic properties by leveraging the excitability and influence of axons that have been spared from injury. Due to the relatively fast improvements in locomotor functions observed in our study (observed within 3-weeks after AAVrg injection), it is likely the therapeutic mechanisms of action are explained by the effects of AAVrg on spared, rather than regenerating axons.

One of the most peculiar findings in our work is the loss of RFP fluorescence observed in the PTEN-KO mice. Prior reports using PTEN-KO have utilized ubiquitous promoters such as the CMV promoter or have utilized transgenic mice which express a reporter gene when exposed to the virally delivered Cre recombinase. While these models apparently exert a more stable reporter gene expression, detecting a down-regulation of the reporter gene while under the hSyn1 promoter is of both extreme interest and concern. A loss of RFP induced by silencing of the hSyn1 promoter may prove to be consistent with an emerging theory on axon regeneration. Specifically, retrograde signals from developed synapses may suppress axon growth, which suggests a need to induce synaptic instability in order to facilitate axon regeneration (26–30). There is little argument that PTEN inhibition induces a regenerative state within affected neurons (2, 8–14, 18, 20, 21, 31, 34–38). Therefore, the suppression of mature synaptic proteins such as synapsin 1 after PTEN-KO would be consistent with the theory of synaptic silencing during regeneration. While silencing of the hSyn1 promoter after PTEN-KO remains speculative, the implications of such findings can be extrapolated to the larger concerns with gene-therapy approaches.

As gene therapies emerge for use in regenerative medicine it will be vital to regulate the effects in a controlled and predictable manner. Use of the hSyn1 promoter may therefore not be the best option for driving genetically induced regeneration, particularly if that growth effect will inevitably become downregulated as the neuron enters a regenerative state. Future work should identify if this same interaction exists using other neuronal specific promoters. One element of gene therapy that also needs to be addressed, particularly after potentially observing toxicity to neurons in the motor cortex, is the long-term repercussions of sustaining a growth and regenerative state in neurons. Prior work has not established a causal link between PTEN-KO and neuronal toxicity, therefore our findings raise concerns about the feasibility of maintaining neurons in a prolonged regenerative state (9, 20).

Previous studies have induced PTEN-KO for sustained periods of time without reporting toxicity or a loss of CST labeling. Specifically, Liu and colleagues (2010) performed PTEN-KO within the motor cortex in neonatal mice and delayed the SCI until 5-months of age (14). Du and colleagues (2015) sustained mice for up to 1 year post-PTEN-KO from the motor cortex without observing toxicity (9). For reasons unknown, our findings deviate from these prior reports and may suggest that either 1) limiting the effects of PTEN-KO to neurons exerts divergent long-term responses compared to using a ubiquitous promoter that may involve astrocytes and microglia in the vicinity of the neuronal soma, or 2) there is something different about using AAVrg’s to knockout PTEN that results in reduced long-term viability.

It is not inconceivable to suggest that a sustained growth response could exert adverse effects at both the level of circuitry but also at the level of neurobiology. One of the main downstream targets that is activated after PTEN-KO, or PTEN inhibition in general, is the hyperactivation of mTOR. mTOR functions as a master regulator of various biological processes, one of which includes autophagy (3, 39–41). When mTOR is active, such as in a PTEN-KO condition, autophagic responses are suppressed (39–41). In several neurodegenerative conditions such as Alzheimer’s, Parkinson’s, or Huntington’s disease, impaired autophagy is linked to the accumulation of neurotoxic protein aggregates and oxidized lipids such as lipofuscin (41–43). Piecing together our findings, it is not implausible that sustaining a prolonged growth response could exert adverse consequences related to the inhibitory effects of mTOR on autophagy. Indeed, an emerging form of cell death termed karyoptosis is induced by chronic inhibition of autophagy (44). Chronic suppression of autophagy and/or induction of karyoptosis after PTEN-KO could explain both a loss of neurons in the motor cortex, as well as the progressive loss in functional abilities observed in our chronic PTEN-KO mice. Taken together, the potential for toxicity caused by long-term sustained growth may have revealed a novel neurobiological barrier for axon regeneration in chronic SCI.

While using AAVrg’s to treat SCI holds significant translational promise, more work is needed to refine the approaches and tease out potential complications associated with this methodology. More importantly, however, our findings have raised a potentially important consideration requiring immediate attention.

Specifically, more knowledge is needed to understand the long-term consequences of sustaining growth in neurons after SCI to both determine if our findings of neuron toxicity are reproducible and to determine if stimulating regeneration using different approaches causes similar complications. Regenerating damaged axons in the human condition will likely require significantly more time compared to rodents due to the difference in the spinal cord size, however, we need to know if neurobiology can sustain the kind of prolonged growth required to induce regeneration over an extended period of time.

Finally, while not observing regeneration of the CST after PTEN-KO may at first seem inconstant with prior literature, our findings are in line with what has been described previously. The lesions in our animals, although remarkably small relative to a clinically relevant contusion lesion, are wider in thickness than prior reports. Here we induced crush injury with forceps approximating 0.2 mm in diameter. Prior studies have utilized forceps that have been filed to be extremely thin, and only when a crush has been performed using extremely thin forceps does axon regeneration of the CST appear to occur (14, 21). This has been suggested to be causal to the ability of astroglia bridges to form and span the ultra-thin lesions, which does not sufficiently occur when the lesions are too wide (21) as in the current study. Zukor and colleagues (2013) described the failure of axon regeneration when astroglia bridges do not form due to experimental lesions induced using 0.5 mm in thickness forceps compared to 0.1 mm thick forceps (21). In our tissue, we did not often see astroglia bridges span the lesion, and when astrocyte bridges were observed they were observed in far-lateral sections or far ventral regions in the cord, both of which are relatively far distances away from the main CST bundle.

Not finding regeneration of the CST even in our acute experiment, while not as hypothesized, is consistent with previous reports considering the nature of our lesions. Our results support a growing idea that astrocytes are indeed essential to support axon regeneration through a lesion and that other inhibitory elements within the lesion mediate regenerative failure (21).

### 3.1 Lay summary of findings

A primary cause for the loss of function experienced after a spinal cord injury (SCI) is due to the disruption of communication to and from the brain. In an uninjured spinal cord, the processes of neurons that carry a signal to other neurons, called axons, project long distances that lead from the brain to the lowest regions of the spinal cord, or from the periphery/lower spinal cord up into the brain. Singular axon processes from neurons within the brain travel uninterrupted down the spinal cord to make connections to other neurons within the spinal cord. Those axons become severed at the location of the injury, preventing communication with the intended target. One of the goals of regeneration after SCI is to force those damaged axons to elongate and grow through the lesion and ideally make connections back to the original and intended target. To date, the ability to induce long-distance regeneration is limited. Regeneration is prevented by inhibitory molecules within the spinal cord, a lack of growth permissive signals/environment, as well as inhibitory mechanisms ongoing within the neurons themselves. One inhibitory molecule that exists inside neurons and actively suppresses regeneration is a protein called the phosphatase and tensin homolog protein (PTEN)(14). Prior work has knocked out PTEN from the genomes of mouse neurons and found that when PTEN is deleted, some axons within the spinal cord are capable of limited regeneration (9, 14).

It is important to note a few things regarding the regenerative observations made in prior studies. First, prior experiments that knocked out PTEN as a growth-promoting strategy have evaluated regeneration only in a single spinal tract at a time within the spinal cord, including the corticospinal tract (CST) or rubrospinal tract (8–11, 14). The CST is both essential to produce voluntary dexterous movements but also has a history of being difficult to coerce to regenerate. Observing regeneration of the CST after PTEN knockout is a huge success, however, relative to all descending CST axons, relatively few grow through the spinal cord lesion after PTEN deletion. While the CST is a singular spinal tract that is important for voluntary movement, many other spinal tracts convey other meaningful functions which ultimately should also regenerate when the goal is to restore normal to near-normal functions. To give a singular example, neurons in a different region of the brain called the Barrington’s nucleus (within the brainstem in a location called the Pons) process information and send axons down the spinal cord to regulate bowel and bladder control which are functions that are not affected if only the CST is regenerated.

There is a need to expand regenerative efforts to include more spinal tracts to maximize the restoration of functions. To date, methods to induce regeneration by inhibiting PTEN have been induced either by using gene-therapy approaches which involve the injection of DNA-delivering viral vectors into the locations in the brain that derive the CST (or other singular fiber tracts), or by delivering a drug systemically that blocks PTEN from functioning (36, 45, 46); both approaches have strengths and limitations. Several of the locations in the brain and brainstem that make up other spinal tracts besides the CST would either be impossible to reach by way of injecting a viral vector, or not advisable due to the potential for off-target effects on surrounding neurons that are not implicated in SCI pathology. Further, it is likely impractical to inject into every location where neurons exist that project into the spinal cord due to the wide distribution throughout the nervous system.

Using systemically delivered drugs to induce regeneration may be capable of reaching a greater diversity of spinal-projecting neurons but may also come with off-target effects, including affecting non-neural cells or neural cells not involved in SCI pathology. Ultimately, we do not know how long it will take for a damaged axon to grow and regenerate in a human spinal cord, and it will likely require sustaining growth for long periods of time. The safety profile for chronically delivering pro-growth compounds systemically remains unknown.

Our project made use of a specific form of a viral vector that is commonly used for gene therapies in both humans and animals. Specifically, a viral particle called an adeno-associated virus (AAV) has been engineered to be taken up by axons at distances far away from the cell body and transported into the cell body (a process called retrograde transport) where it can elicit the desired genetic effects (17). Importantly, AAVs that are used for this experimental purpose cannot replicate so when injected into the spinal cord, only cells or axon processes local to the injection site take up the virus. Our study took advantage of this retrograde property of the novel AAV and through injection directly above (rostral) to the lesion site, we attempted to knockout PTEN from most of the spinal projecting neurons throughout the neuroaxis without affecting many off-target neurons.

We delivered our gene therapy in both acute and chronic SCI conditions and observed that several mice with a near-complete loss of function regained weight supporting abilities regardless of when the AAV was delivered. In mice that were treated immediately after injury, we saw a corresponding increase in axon growth into the lesions. In mice that were treated in chronic SCI, we observed that the worse a mouse was able to function at the time of treatment, the better that mouse performed after knocking out PTEN. However, we also observed that some mice that could weight support before treatment, lost weight supporting abilities after knocking out PTEN. Most importantly, in the mice that responded with a gain of function after knocking out PTEN, the functional improvements were not retained for longer than 9-weeks.

We kept chronically treated mice alive for 6-months after treatment to monitor for motor abilities and found that all mice receiving PTEN knockout lost deletion-mediated improvements from 9- to 33-weeks post-treatment. When evaluating tissue obtained from these mice, we detected a possible toxic response to neurons in the motor cortex that give rise to the CST. We currently do not know the mechanisms underlying toxicity caused by sustained PTEN knockout. At this time, our experimental findings cannot be directly translated into humans for several reasons. First, our data raise concern for the long-term consequences of sustaining growth through PTEN interference, therefore further work is needed to identify the safety of similar approaches and optimize the delivery methods. Next, our study made use of mice which have been genetically engineered to be capable of responding to the gene therapy used in this study. More sophisticated approaches will be required to create a similar gene-therapy approach that will function in other animals including humans (13). Finally, our observation of injury-severity dependent responses to PTEN knockout suggests that more work is needed to determine what injury characteristics (complete vs incomplete, and what fiber tracts should or should not be spared from injury) will benefit from retrogradely transported AAVs to elicit axon growth. While observing any improvement in function in chronic SCI conditions is exciting, it remains important to interpret our findings as yet another small step forward. There is still a significant amount of work ahead before an analogous treatment similar to what was performed in this study could be considered for experimental purposes in humans.

## 4.0 Conclusions

To conclude, use of AAVrg’s to knockout PTEN exerts dynamic effects that are dependent upon the time applied post-SCI, the injury severity, and the duration of recovery post-treatment. PTEN-KO using AAVrg’s improves locomotor abilities in near-complete SCI when used to treat both acute and chronic SCI, but the effects on less-severe injuries appear detrimental to weight-supporting abilities. We observed that long-term PTEN deletion showed negative effects even though there was an initial improvement in the most severe injuries. While the use of spinal-injections of AAVrg’s provides a promising approach to target the breadth of axons implicated in the pathophysiology of SCI, future work is required to optimize the methods and delivery approaches, as well as to identify the injury conditions most likely to experience a therapeutic gain.

## 5.0 Methods

### 5.1 Animals and Spinal Cord Injury Modelling

All procedures were approved by the University of Kentucky Institutional Animal Care and use Committee. C57BL/6J mice with loxp sites flanking the catalytic domain of PTEN (B6.129S4-PTEN^tm1hwu^/j; strain #006440; The Jackson Laboratory) were obtained and bread as homozygous breading pairs. Male mice were used for acute behavioral studies while female mice were used for chronic behavioral studies. Tract tracing experiments for the acute-treatment time point utilized a mix of male and female mice. Due to the long-timepoint used in the chronic experiment, female mice were used to mitigate unexpected mortality and health issues that arise in male mice (47). Experimental groups and group sizes are included in Table 1. A graphical overview of experimental methods and timelines is available in Figure 1.

Mice were injured between 11-13 weeks of age through a complete spinal crush injury at the T9 vertebral spinal level. Following dissection of the skin and musculature above the T9 vertebral process, a T9 laminectomy was performed and then crush injuries were performed using fine-tipped forceps as described in prior literature (9, 14). Forceps were closed on the spinal cord using care to ensure the tips were scraping the ventral aspect of the vertebral column to minimize the risk of sparing axons at the lesion. Significant force was applied to the forceps and the forceps were held closed for 8-seconds. For acute experiments, mice were suspended by vertebral clips immediately after crush SCI and 2-µL of AAVrg was injected at 2-mm rostral to the lesion, centered on midline of the spinal cord, and approximately 0.7-mm below the dura surface. Injections were performed on a stereotactic device using a Hamilton syringe containing a leur lock tip for interchangeable tips (80001; Hamilton Company, Reeno, NV). Fine glass-pulled pipette tips were purchased at a calibrated 10.0 µm diameter (TIP10TW1-L; World Precision Instruments, Sarasto, FL). The entire Hamilton syringe with fitted pipette tip was filled with mineral oil to create a hydraulic continuity between the plunger and aspirated virus. Virus was injected at a rate of 0.3 µL/minute and the needle was left in place for 3 minutes after finishing the injection. For chronic experiments, the spinal cord was re-exposed at 12 weeks post-SCI and 2 µL of AAVrg was injected as just described.

For all surgical procedures, mice were anesthetized using intraperitoneal injections of ketamine (100.0 mg/kg) and xylazine (10.0 mg/kg). Incisions of mice were sutured using absorbable sutures and the skin was closed using non-absorbable sutures. Mice were given antibiotic (enrofloxacin; 5.0 mg/kg), and saline (1.0- mL/day) support for 5 consecutive days after SCI. Analgesic support was provided using a 1x delivery of buprenorphine slow-release formulation (Buprenex SR, 1.0 mg/kg). Bladders of mice were expressed 2x/day for the duration of the study consisting of 9 weeks for acute experiments and 9 months for chronic experiments. AAVrg’s were purchased from Addgene and included either AAVrg-hSyn-Cre-p2A-dTomato (1.5x10^13^ vg/mL; 107738-AAVrg; Addgene, Watertown, MA), which was used to knockout PTEN, or AAVrg-hSyn-mCherry (2.0x10^13^ vg/mL; 114472-AAVrg; Addgene, Watertown, MA) as a vector control. For our initial co-labeling experiments to identify the distribution of transfected neurons throughout the brain, we used AAVrg-hSyn-eGFP (1.7x10^13^ vg/mL; 50465-AAVrg; Addgene, Watertown, MA). AAV’s were used undiluted after 2 freeze-thaw cycles, one to allocate and the other prior to injections, as recommended by the manufacturer. pAAV-hSyn-Cre-P2A-dTomato was a gift from Rylan Larsen (Addgene viral prep # 107738-AAVrg; http://n2t.net/addgene:107738; RRID:Addgene_107738). pAAV-hSyn-mCherry was a gift from Karl Deisseroth (Addgene viral prep # 114472-AAVrg; http://n2t.net/addgene:114472; RRID:Addgene_114472). pAAV-hSyn-eGFP was a gift from Bryan Roth (Addgene viral prep # 50465-AAVrg; http://n2t.net/addgene:50465; RRID:Addgene_50465).

Mice in acute experiments were allowed to survive for up to 9 weeks post-SCI. The 9-week time-point was chosen due to prior work demonstrating a need to prolong experiments at least 9 weeks in order to detect regeneration of the corticospinal tract (CST) in mice receiving PTEN-KO delivered to the sensory-motor cortex (14). For chronic experiments, mice were allowed to survive for 9 months. The 9-month time-point was similarly chosen due to the singular report of PTEN-KO performed in chronic SCI that identified limited axon regeneration of the corticospinal tract at 4 months post-knockout with better regenerative effects detected at 7 months (9). For chronic experiments, mice survived for a total of 3 months before treatment, and 6 months post-treatment, with the 6-month post-treatment time point being determined by the emergence of severe autophagia in several mice, unrelated to group assignment, causing early termination of the study.

### 5.1 Tract Tracing Procedures

Experiments using tract tracing procedures utilized either Fluorogold (1.0% w/v; Fluorochrome; Denver, CO) injections into the spinal cord rostral to the lesion as a retrograde tracer or injections of biotinylated dextran amines (BDA, D1956; 10,000 kD, 10.0% w/v, ThermoFisher Scientific, Waltham, MA) into the motor cortex as an anterograde tracer to visualize the CST. Tracer injections were performed two weeks prior to euthanasia. Mice were anesthetized again with ketamine (100.0 mg/kg) and xylazine (10.0 mg/kg) and either had their lesion sites re-exposed or received craniotomies to expose the motor cortex of mice. Craniotomies were performed by gentle grinding of the skull using a rotary tool, and removal of the bone over the injection coordinates at -0.3, -0.8, -1.3 mm caudal to bregma at all 1.5, 1.0, -1.0, and -1.5 mm lateral to bregma. BDA injections (0.4 μL/injection) were made into all 12 locations at a rate of 0.3 μL/min using pre-pulled glass pipettes as described above, at a depth of approximately 0.6 mm below the dura surface. The needle was left in place for at least 1 minute after each injection. For Fluorogold injections, the spinal cord was revealed after removal of developed scar tissue over the dura and a pulled glass pipette, prepared as described above, was lowered approximately 0.7 mm below the dura surface. 1.0-μL of Fluorogold was injected into the spinal cord at least 2.0 mm rostral to the lesion to ensure effective labeling of axons that have died back away from the site of injury. The needle was left in place for at least 3-minutes after injection. Skin lesions over the skull were closed using reflex clips, while lesions over the spine were closed using non-absorbable sutures. All mice were treated with Buprenorphine-SR for analgesic support. For tract-tracing of the acute experiments only 1 mouse in the control group survived until scheduled euthanasia while 4 mice in the PTEN-KO survived. While not ideal, the main question under investigation was about the survival of neurons in the PTEN-KO group and viability of neurons labeled with the AAVrg was apparent in control-treated mice. In the chronic experiment, 3 control mice and 3 PTEN-KO mice survived.

### 5.3 Behavioral Monitoring

Due to the severe nature of complete crush SCI models, we did not predict any mouse to recover enough function to perform weight-supported stepping. Instead, if improvements were to be made, we estimated kinematic improvements would be detectable through increased range of motion of legs. We chose to utilize the Basso Beattie and Bresnahan scale of locomotor improvements (22) (BBB), which was developed for rats with contusive SCI, rather than the Basso Mouse Scale (23) (BMS) that is developed for mice with contusive SCI, due to more resolution within the BBB at lower ranges of function. Unfortunately, there are no other standardized and objective measures within the SCI field to evaluate hind limb kinematic improvements prior to weight-supported abilities, which underlies the strength of using the BBB scale to assess this range on the continuum of recovery.

As described by Basso and colleagues (2006), detection of range of motion of all joints in mice is challenging which is the rationale for only assessing ankle movement on the BMS as opposed to assessing the ankle, knee, and hip in the BBB. However, we chose to accept this limitation and move forward with utilizing the BBB rating scale. To acquire open field scores, mice were placed in an unfilled children’s pool and allowed to explore for 4-minutes. Raters who were blinded to experimental groups assessed the hind-limb functions of mice using the 21-point BBB scale. Early stages of the BBB scale assess the range of motion of each of the 3 joints of the hind limbs indicated as either no movement, slight movement, or extensive movement, and include observations to characterize sweeping behaviors, all which emerge before plantar placement of the paws.

Evaluating the range of motion of all joints, characterizing sweeping behaviors, and separating pre- (BBB scores of 8 and below) vs post-weight supporting abilities (BBB scores above an 8) marks an increase in resolution on the BBB scale relative to the BMS scale for severe conditions of SCI. The BMS scale only evaluates the ankle joint, doesn’t characterize sweeping behaviors, and combines pre-weight supported plantar placement, weight supported stance, and up to consistent weight-supported dorsal stepping as a single score (score of 3 on the BMS). The clustering of kinematic behaviors that transition from pre- to post-weight supporting abilities in the BMS scale provides significantly less resolution for discriminating differences occurring in severe, near-complete SCI. Mice were assessed at 1-week post-SCI and bi-weekly thereafter until euthanasia.

### 5.4 Immunohistochemistry

#### 5.4.1 Tissue Prep

At 8-weeks post-SCI mice were again anesthetized using an overdose of ketamine (∼200.0 mg/kg) and xylazine (∼20.0 mg/kg) and perfused using 0.1 M phosphate buffered saline (PBS) followed by perfusion with 4.0% formaldehyde prepared from prilled paraformaldehyde (441244; Sigma-Aldrich) in PBS. The brains and spinal cords of mice were isolated and post-fixed at room temperature in 4.0% formaldehyde for 2 hours. After post-fixation, cords were transferred and acclimated to 30.0% sucrose over the next week. The spinal cords and brains were frozen in Shandon M1 Embedding Matrix (FIS1310; ThermoFisher Scientific) and cut sagittal on a cryostat at -15.0 °C at 20.0 µm thickness. Cords were blocked together with at least two mice per group embedded in each block. For each immunohistochemistry stain, one section every 140.0 µm of tissue thickness was assessed. Brains were blocked together with one mouse per group embedded in each block.

The brains were cut between the caudal end of the cerebellum and the rostral aspect of the motor cortex. Brains were cut at 20-30 µm and captured as serial sections on slides.

#### 5.4.2 Immunohistochemistry Sample Processing

Before performing immunohistochemistry (IHC), sections were dehydrated by immersion in graded ethanol baths containing 70-, 95-, and 100% ethanol and cleared of lipids by immersion in xylene for 5-minutes. We have found that clearing sections using organic solvents can often improve antibody penetration and labeling quality of thicker sections, particularly in the white matter, when IHC labeling is performed on slides. After rehydration of sections, when performing chromogen labeling, slides were treated with quenching buffer consisting of 60/40% PBS/methanol and 0.3% H_2_O_2_ for 30-minues then washed in PBS. When performing fluorescent labeling, or after quenching during chromogen labeling, unspecific labeling was blocked using 60-minute incubations in 5.0% normal goat serum solution made in PBS and 0.1% Triton-X 100. All primary antibody labeling was performed at room temperature overnight. Secondary antibodies include Alexafluor-conjugated Goat anti-IgG/IgY (1:500; A11008, A21449; ThermoFisher Scientific.), peroxidase-conjugated Goat anti-IgG (1:250; PI-1000, Vector Laboratories), or alkaline phosphatase-conjugated goat anti-IgG/IgY (1:1000; A16057; ThermoFisher Scientific) and were incubated for 1-hour in PBS and 0.1% Triton-X 100 at room temperature. Spinal cord sections labeled for fluorescent IHC were treated with True Black (0.5x; 23007; Biotium) after antibody labeling to reduce auto-fluorescence caused by lipofuscin accumulation in macrophages.

#### 5.4.3 Spinal Cord Labeling

Sections were labeled to visualize the astrocyte scar using antibodies that target glial fibrillary acidic protein (GFAP; 1:3000; Aves Labs) and co-labeled with β-tubulin III (β3-Tub; 1:2000, 5568S; Cell Signaling Technologies) to visualize axons within the lesion. The red fluorescence from both mCherry and dTomato, when present (see discussion below), was bright enough to not require immunohistochemical labeling.

Serotonin (5-HT) positive axons were identified using chromogen labeling against 5-HT (1:4000; 20080; ImmunoStar) and were either not co-labeled in the first acute experiment, or were co-labeled with GFAP (1:5000) using alkaline phosphatase-conjugated secondaries to visualize the astroglia borders. The CST was labeled using the Avidin Biotin Complex method (ABC) to detect the BDA by incubating in ABC (5 µL/mL A and B; PK-6100; Vector Laboratories) overnight at room temperature and co-labeled against GFAP (1:5000) to visualize astroglia borders.

#### 5.4.4 Brain Labeling

Brain sections were examined to identify populations of neurons affected by the AAVrg approach. Anatomical identification of the motor cortex, red nucleus, caudal Raphe nucleus, Locus Coeruleus, and Barrington’s nucleus were made either through landmark identification, morphology, or histological verification. The Locus Coeruleus was identified using histological identification of dopamine beta-hydroxylase (DBH; 1:2000; 22806; ImmunoStar) positive neuron clusters at the trunk of the cerebellar peduncle shaped as a half moon, while the Barrington’s nucleus was identified as DBH^-^ neurons clusters cradled by the half-moon of the Locus Coeruleus (48). Serotonergic neurons within the caudal Raphe nucleus were histologically identified by labeling against 5-HT (1:4000) and detected in the ventral-medial aspect of the brainstem. The red nucleus is easily identifiable by a bi-lateral circular cluster of neurons within the mid-brain that label vividly using AAVrg approaches, while neurons of the motor cortex are identifiable as the only neurons labeled within layer 5 of the cortex. In animals labeled with Fluorogold, similar brain regions were assessed for the expression of both Fluorogold and the RFP expressed by the viral genome to determine differences in viability of infected neurons between groups.

#### 5.4.5 Imaging and Quantification

Fluorescently labeled spinal cord sections were imaged using confocal microscopy to obtain the necessary pixel resolution and signal-to-noise ratio required for evaluating fine axon fibers within the lesion. Brain sections were imaged using conventional epifluorescence. Z-stack images were obtained and compressed into a single maximal intensity image projection file for analysis. GFAP lesion borders were identified on spinal cord sections and traced using Halo software (Indica Labs, Albuquerque, NM). The percentage and intensity of the lesion filled with axonal labeling was obtained after setting threshold parameters to identify positively labeled pixels. To quantify 5-HT labeling, first, sections were evaluated for evidence of axon growth across the lesion. The caudal lesion boundary of sections with evidence of 5-HT labeling was traced and an outward duplication of the trace performed to make new boundaries every 25- or 50-µm from the lesion border for the acute and chronic experiments respectively. Every location where a 5-HT axon crossed a trace was identified and counted. Due to the often beaded and non-continuous appearance of 5-HT labeling, axons with a visible trajectory crossing or approximating a boundary were also counted to ensure an accurate representation of axon growth. The frequency of axon-trace interactions was quantified and compared across groups. The average amount of axons crossing at each assessed distance per section was used for analysis.

For BDA labeling of the CST there were no BDA labeled axons caudal to the lesion in any animal, therefore quantification of regenerated CST was not performed. Instead, due to an unexpected but apparent loss of CST labeling in chronically treated PTEN-KO mice, the quality of CST labeling was categorically coded as either completely filled, moderately filled, sparse labeling, or no labeling. A reviewer who was blinded to group conditions analyzed all labeled sections for the visibility of CST labeling. Completely filled CST labeling was apparent through a dense central CST bundle usually apparent in multiple midline sections. Moderately filled CST labeling was considered when a singular CST bundle could be identified but had less density or appeared on fewer sections compared to completely filled examples. Sparse labeling was considered when axon collaterals could be detected in the grey matter, but no sections contained a singular or identifiable CST bundle. No labeling was considered when no axon collaterals were observed. Due to the technical complexity of BDA cortical injections, any animal with surgical notes that conferred question about the labeling quality were excluded from assessment. Mice with evidence of complete or moderately labeled CST bundles were analyzed in one group, and mice with sparse or no labeling were delegated into a second group. The frequency of complete/moderately labeled vs sparse/no labeling were assessed using Chi-Squared.

Brain sections were labeled using the same immunohistochemical strategy as for spinal cord sections. Neuronal populations of interest were identified to qualitatively assess the distribution of affected brain regions at 2-weeks, 9-weeks, or 9-months post-SCI. Prior work by Blackmore and colleagues (2018) had identified minimal co-labeling within the Raphe nucleus and almost no labeling within the Locus Coeruleus at 2-weeks post-treatment (24). Our study evaluated neurons at much longer time points post-treatment which gives significantly more time to either stop expressing, or start expressing, in ways undescribed from prior literature.

### 5.5 Statistics

Repeated-measures Analysis of Variance (ANOVA) was used to assess BBB scores as well as 5-HT axon regeneration beyond the lesion. Post-hoc comparisons were performed using Fisher’s least significant differences when evaluating between-group effects on the repeated measures model, and Dunnett’s pairwise comparisons when comparing within group effects on repeated measures designs. T-tests were used to compare axon labeling within the lesions, and Chi-Squared was used to evaluate the frequency of occurrences when data was categorical. Probability values obtained at *p* < 0.05 were considered significant.

## Data Sharing

All data from this manuscript will be publicly available through the Open Data Commons for Spinal Cord Injury (www.odc-sci.org).

## Authorship Contributions

**Conceptualization**: A.S. & J.G. **Methodology**: A.S., R.K., W.B., G.H., E.G., O.W. **Validation**: A.S., J.G. **Formal Analyses**: A.S., J.G. **Investigation**: A.S., J.G. **Data Curation**: A.S., J.G. **Writing**: A.S., J.G. **Project Administration**: A.S., J.G. **Resources**: J.G. **Funding Acquisition**: A.S., J.G. **Supervision**: J.G.

## Abbreviations

5-HT: serotonin
AAV: adeno-associated virus
AAVrg: retrograde psuedotyped adeno associated virus
ABC: avidin biotin complex
ANOVA: analysis of variance
β3-Tub: beta III tubulin
BBB: Basso Beattie and Bresnahan scale of locomotor recovery
BDA: biotinylated dextran amine
BMS: Basso Mouse Scale
CNS: central nervous system
CST: corticospinal tract
DBH: dopamine beta hydroxylase
eGFP: enhanced green fluorescent protein
GFAP: glial fibrillary acidic protein
hSyn1: human synapsin 1 promoter
IHC: immunohistochemistry
KO: genetic knockout
mTOR: mammalian target of rapamycin
PBS: phosphate buffered saline
PTEN: phosphatase and tensin homolog protein
RFP: red fluorescent protein
SCI: spinal cord injury.

## Acknowledgements

Thank you to the Wings for Life Foundation for providing funding to support this project. Thank you to Taylor Valentino for facilitating pilot experiments for this project. Thank you to Michael Leighton for your help as an SCI lived experience advocate for helping construct the layman’s abstract. Confocal microscopy was performed at the University of Kentucky’s Light Microscopy Core. pAAV-hSyn-Cre-P2A-dTomato was a gift from Rylan Larsen (Addgene viral prep # 107738-AAVrg; http://n2t.net/addgene:107738; RRID:Addgene_107738). pAAV-hSyn-mCherry was a gift from Karl Deisseroth (Addgene viral prep # 114472-AAVrg; http://n2t.net/addgene:114472; RRID:Addgene_114472). pAAV-hSyn-eGFP was a gift from Bryan Roth (Addgene viral prep # 50465-AAVrg; http://n2t.net/addgene:50465; RRID:Addgene_50465).

---

**Supplemental Figure 1.**
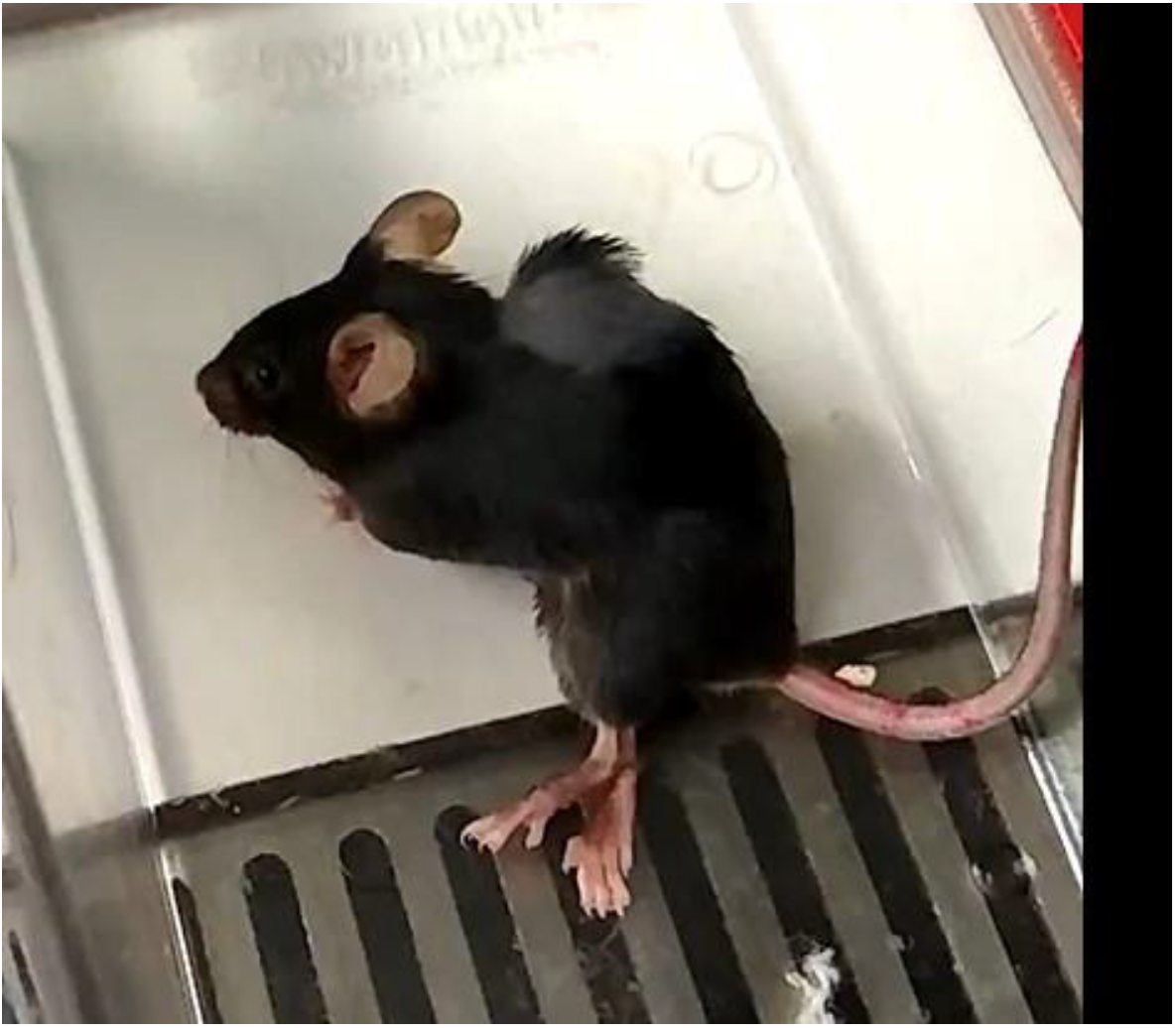
PTEN-KO using AAVrg resulted in sustained contraction of the abdomen. Mice receiving mid-thoracic spinal injections of AAVrg’s to knockout PTEN experienced observable sustained contraction of the abdomen. The extent to which increased abdominal tone and contraction affected weight supporting abilities or trunk stability remains unknown.

