## Supplementary figures and images for "PTEN knockout using retrogradely transported AAVs restores locomotor abilities in both acute and chronic spinal cord injury"

### Supplemental Figure 1

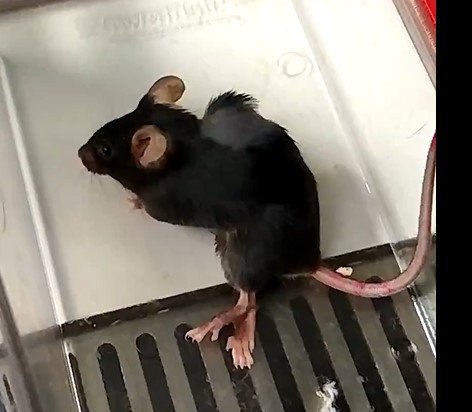
